# The first next-generation sequencing approach to the mitochondrial phylogeny of African monogenean parasites (Platyhelminthes: Gyrodactylidae and Dactylogyridae)

**DOI:** 10.1101/283788

**Authors:** Maarten P.M. Vanhove, Andrew G. Briscoe, Michiel W.P. Jorissen, D. Tim J. Littlewood, Tine Huyse

## Abstract

**Background:** Monogenean flatworms are the main ectoparasites of fishes. Representatives of the species-rich families Gyrodactylidae and Dactylogyridae, especially those infecting cichlid fishes and clariid catfishes, are important parasites in African aquaculture, even more so due to the massive anthropogenic translocation of their hosts worldwide. Several questions on their evolution, such as the phylogenetic position of *Macrogyrodactylus* and the highly speciose *Gyrodactylus*, remain unresolved with available molecular markers. Also, diagnostics and population-level research would benefit from the development of higher-resolution genetic markers. We aim to advance genetic work on African monogeneans by providing mitogenomic data of four species (two each belonging to the Gyrodactylidae and Dactylogyridae), and analysing their gene sequences and gene order from a phylogenetic perspective.

**Results:** Based on Illumina technology, the first four mitochondrial genomes of African monogeneans were assembled and annotated for the cichlid parasites *Gyrodactylus nyanzae*, *Cichlidogyrus halli*, *Cichlidogyrus mbirizei* (near-complete mitogenome) and the catfish parasite *Macrogyrodactylus karibae* (near-complete mitogenome). The start codon TTG is new for *Gyrodactylus* and for the Dactylogyridae, as is the incomplete stop codon TA for the Dactylogyridae. The most variable markers are *nad* genes and these are under relaxed selection. Especially *nad*2 is promising for primer development. Gene order was identical for protein-coding genes and differed between the African representatives of these families only in a tRNA gene transposition. A mitochondrial phylogeny based on an alignment of nearly 12,500 bp including 12 protein-coding and two ribosomal RNA genes confirms that the Neotropical oviparous *Aglaiogyrodactylus forficulatus* takes a sister group position with respect to the other gyrodactylids, rather than the supposedly ‘primitive’ African *Macrogyrodactylus*. Inclusion of the African *Gyrodactylus nyanzae* confirms the paraphyly of *Gyrodactylus*. The position of the African dactylogyrid *Cichlidogyrus* is unresolved, although gene order suggests it is closely related to marine ancyrocephalines.

**Conclusions:** The amount of mitogenomic data available for gyrodactylids and dactylogyrids is increased by roughly one-third. Our study underscores the potential of mitochondrial genes and gene order in flatworm phylogenetics, and of next-generation sequencing for marker development for these non-model helminths for which few primers are available while they constitute a risk to tropical aquaculture.

## Background

Ectoparasitic infections in bony fishes are dominated by monogeneans (Cribb et al. 2002). Among their most species-rich taxa are the Gyrodactylidae and the Dactylogyridae (Pugachev et al. 2009). These include, respectively, the supergenera *Dactylogyrus* and *Gyrodactylus*, some of the most significant radiations of flatworm fish parasites (Cribb et al., 2002). Around 500 species of *Gyrodactylus* species have been described at present (Zahradníčková et al. 2016 and references therein), but the estimated species number is much higher (Bakke et al., 2002). These minute flatworms attach to their host by means of a characteristic and diagnostic opisthaptor (Paladini et al. 2017). The resulting disruption of the epidermis can facilitate secondary infections (Bakke et al. 2007). Some genera within these families, such as *Gyrodactylus*, *Macrogyrodactylus*, *Dactylogyrus* and *Cichlidogyrus* include important fish pathogens, causing mortalities especially in captive-reared stocks and after anthropogenic co-introduction outside of their native range (Paperna 1996; Pugachev et al. 2009; Paladini et al. 2017). In Africa, the most important aquaculture fishes are species of the Cichlidae and Clariidae, including respectively the Nile tilapia and the North African catfish, which have been introduced worldwide (Pouomogne 2010; Deines et al. 2016). These fish families are also relatively well-studied for monogenean parasites (e.g. Beletew et al. 2016; Zahradníčková et al. 2016). They harbour several originally African monogeneans that are widely distributed within and outside Africa, and that are important in the study of parasite ecology, evolution and invasion biology because of the economic and scientific importance of their hosts (Vanhove et al. 2016).

In view of the important threats that disease poses to the sustainable development of aquaculture in developing countries, a better monitoring and identification of aquatic pathogens is vital (Bondad-Reantaso et al. 2005). In Africa, more understanding of the diversity and ecology of fish parasites is needed to implement government policies on aquatic health management (Akoll et al. 2012a). Nevertheless, even in the light of the rampant human translocation of fishes and the associated risk of parasite co-introductions, the lack of monitoring is clear (e.g. Smit et al. 2017). Monogeneans, in particular, have been assessed as high-risk parasites in African aquaculture (Akoll et al. 2012b). Since common procedures for the identification of these monogeneans are lethal to the host and require a high level of technical expertise, non-intrusive molecular diagnostics are called for (e.g. Ek-Huchim et al. 2012 for *Cichlidogyrus*). However, there is a lack of high-resolution molecular markers for these animals (Vanhove et al. 2016).

In addition, the phylogenetic position of African monogenean lineages, including several endemic or recently discovered genera, is often poorly understood. For example, the currently most frequently used markers, situated in the nuclear ribosomal DNA region, have not yet fully resolved the position of the typically African *Macrogyrodactylus*. This genus’ representatives infect clariid catfishes, among other hosts (Barson et al. 2010; Přikrylová et al. 2013). Malmberg (1998) suggested, based on morphological data, that the genus comprises the earliest divergent lineage of gyrodactylids. This is a family of mainly viviparous monogeneans, although with some oviparous representatives (Bakke et al. 2007). However, Malmberg’s hypothesis was recently contradicted by mitogenomic phylogenetics where the Neotropical oviparous gyrodactylid *Aglaiogyrodactylus forficulatus* is sister to all other, viviparous, family members (Bachmann et al. 2016), while nuclear data places *Macrogyrodactylus* instead with other viviparous lineages (Přikrylová et al. 2013). Another long-standing issue in the phylogeny of this monogenean family, is the status of its most speciose and well-studied genus, *Gyrodactylus* (Gilmore et al. 2012 and references therein), first suggested to be paraphyletic by Kritsky & Boeger (2003).

Recently, next-generation sequencing (NGS) approaches have facilitated marker development for non-model helminths (Minárik et al. 2014); this includes the assembly of mitogenomes for fish helminths (Hahn et al. 2013; Brabec et al. 2015). Here we want to apply this approach to the understudied, but highly diverse, African monogenean fauna. We targeted two species of *Cichlidogyrus* (Dactylogyridae), the most speciose monogenean genus infecting African cichlid fishes (Pariselle & Euzet 2009); one gyrodactylid parasite of cichlids; and a representative of *Macrogyrodactylus*. Through phylogenomic and gene order analysis, we address the following questions:

1. Are the Neotropical oviparous gyrodactylids still basal in a mitochondrial phylogeny when including the viviparous *Macrogyrodactylus*, which is supposedly the earliest divergent gyrodactylid lineage according to Malmberg (1998)?
2. Does the phylogeny based on mitogenomic data confirm the paraphyly of *Gyrodactylus*?
3. Do the African representatives of the Gyrodactylidae have the same gene order in their mitochondrial genome as the known Palearctic ones?
4. Do the African freshwater representatives of the Dactylogyridae have the same gene order as seen in the only known dactylogyrid mitogenomes, from a Palearctic freshwater and an Indo-Pacific marine species?

## Methods

### Sampling

Fish hosts were collected in the Haut-Katanga province of the D.R. Congo in 2014. Two individuals of North African catfish *Clarias gariepinus* (vouchers URA 2014-P-1-004 at the Université de Lubumbashi, D.R. Congo, and MRAC 2015-06-P tag AB49120835 at the Royal Museum for Central Africa (RMCA), Belgium) were caught in the Kiswishi River at Futuka Farm (11°29’S 27°39’E) on August 30^th^-31^st^ and a hybrid between Nile tilapia *Oreochromis niloticus* and Mweru tilapia *Oreochromis mweruensis* (voucher MRAC 2015-06-P tag 2655) at the Kipopo station of the Institut National pour l’Etude et la Recherche Agronomiques (11°34’S 27°21’E) on August 27^th^. Hosts were sacrificed using an overdose of tricaine methanesulfonate (MS222). Parasites isolated either *in situ* or later from preserved fish gills were fixed and preserved in analytical-grade ethanol. Individual monogenean specimens were identified on the basis of their morphology using keys and features described in Pariselle & Euzet (2009), Barson et al. (2010) and Zahradníčková et al. (2016). Identified specimens were pooled per species in absolute ethanol: four specimens of *Macrogyrodactylus karibae* (supplemented with two extracts from Barson et al. (2010)), 43 of *Cichlidogyrus mbirizei*, 18 of *Cichlidogyrus halli* and 44 of *Gyrodactylus nyanzae*. While *M. karibae* is a typical gill parasite of *Clarias gariepinus* known from southern Africa (Barson et al. 2010 and references therein), *G. nyanzae* and especially *C. halli* are known from a wide range of cichlids throughout Africa (Pariselle & Euzet 2009; Zahradníčková et al. 2016). The two latter species have previously been reported from tilapias in the Haut-Katanga province (Jorissen et al. 2017). *Cichlidogyrus mbirizei* was only recently described from the Lake Tanganyika endemic *Oreochromis tanganicae* (Muterezi Bukinga et al. 2012). It was afterwards also found on Nile tilapia and its hybrid *O. niloticus* x *mossambicus* (Lerssutthichawal et al. 2016; Lim et al. 2016) and is here for the first time reported from *O. niloticus* x *mweruensis*. The abovementioned reports of both species of *Cichlidogyrus* include co-introductions outside Africa, in nature and in aquaculture settings.

### DNA extraction and sequence assembly

Total genomic DNA was extracted using the DNeasy Blood and Tissue Kit (Qiagen) following the manufacturer’s instructions. The amount of double-stranded DNA isolated was measured with Qubit® 2.0 Fluorometer (Life Technologies, Paisley, UK) yielding 0.9 (*M. karibae*), 3.3 (*C. halli*), 3.2 (*C. mbirizei*) and 1.8 (*G. nyanzae*) ng/μl total DNA.

Samples for NGS were prepared and run at the DNA Sequencing Facility of the Natural History Museum, London, UK. Genomic DNA was indexed and libraries prepared using the TruSeq Nano DNA Sample Preparation Kit (Illumina, Inc., San Diego, USA), and run simultaneously on a MiSeq Illumina sequencer yielding 300 bp long paired-end reads. The new mitogenomes were directly assembled using Geneious v. 8.1.9 (Kearse et al. 2012). The sequences were first trimmed (error probability: 0.05, maximum ambiguity: 1) and then assembled. Partial *cox*1 sequences of *Gyrodactylus salaris* (NC008815 (Huyse et al. 2007)) (for *G. nyanzae*), *Macrogyrodactylus clarii* (GU252718 (Barson et al. 2010)) (for *M. karibae*) and *Cichlidogyrus zambezensis* (KT037411 (Vanhove et al. 2015)) (for representatives of *Cichlidogyrus*) were used as reference sequence to extract *cox*1 reads from the Illumina genomic readpool to form the consensus sequence to subsequently map the reads on successive iterations. Trimmed reads were mapped back to the contigs in order to estimate the full mitochondrial genome coverage, trim the overlapping regions to create a circular molecule, and to inspect for potential mapping/assembly errors in problematic regions such as repetitive regions (Briscoe et al. 2016b).

Using nuclear ribosomal RNA gene sequences for *Cichlidogyrus halli* and *Macrogyrodactylus congolensis* from GenBank (accessions: HE792784 (Mendlová et al. 2012) and HF548680 (Přikrylová et al. 2013) respectively), fragments of the ribosomal RNA operon were identified and assembled using the same iterative process as described for the mitochondrial genome. Exact coding positions of the 18S and 28S nuclear rDNAs, as well as the respective 5′ and 3′ boundaries of the external transcribed spacers, were determined using RNAmmer (Lagesen et al. 2007). Subsequently the complete annotation was compared with the fully-annotated human rDNA repeating unit (GenBank accession: HSU13369).

### Mitogenome annotation

The identity and boundaries of individual protein-coding genes (PCGs) and ribosomal RNA (rRNA) genes were determined using the MITOS web server (Bernt et al. 2013) in combination with the visualisation of open reading frames in Geneious and a comparison with alignments of available mitogenomes of monopisthocotylean monogeneans. In addition to MITOS, the ARWEN v. 1.2 (Laslett & Canbäck 2008) and tRNAscan-SE v. 2.0 (Lowe & Eddy 1997) web servers were used to identify the tRNA-coding regions. When results between applications conflicted, the solution proposing a 7 bp acceptor stem was chosen. We checked for repeat regions with Tandem Repeats Finder (Benson 1999) and YASS (Noe & Kucherov 2005). The resulting mitogenomes were visualised in OGDRAW v. 1.1 (Lohse et al. 2013).

### Alignment, sequence analysis, phylogenetic reconstruction and gene order analysis

Ribosomal RNA genes were aligned by MAFFT v. 7 (Katoh & Standley 2013) using the Q-INS-i iterative refinement method, taking into account RNA secondary structure (Katoh & Toh 2008). Codon-based alignment of all obtained PCGs was performed under the echinoderm and flatworm mitochondrial genetic code (Telford et al. 2000) using MUSCLE (Edgar 2004) implemented in SeaView v. 4.6.2 (Gouy et al. 2010). Since omitting unreliable portions of the alignment may increase resolution in phylogenomic reconstructions (Philippe et al. 2017), an alternative alignment was obtained by trimming in Gblocks v. 0.91b (Castresana 2000), implemented for the PCGs in TranslatorX (Abascal et al. 2010), carrying out codon-based MAFFT alignment followed by alignment cleaning in Gblocks. Options for a less stringent selection were selected, allowing smaller final blocks, gap positions within the final blocks, and less strict flanking positions. Especially for smaller datasets, trimming entails the risk of removing information contributing to phylogenetic signal (Philippe et al. 2017). Therefore, likelihood mapping (Strimmer & von Haeseler 1997) was performed in TREE-PUZZLE v. 5.3 (Schmidt et al. 2002) to compare the phylogenetic content of the complete and trimmed concatenated alignment. The percentage of fully, partially and unresolved quartets was 99.4, 0.5 and 0.1 in both cases, hence trimming did not increase phylogenetic content and the original alignment was preferred for downstream analyses (Fig. 1). Comparing, in DAMBE (Xia 2017), the index of substitution saturation with its critical value at which sequences would start to fail to recover the true phylogeny, indicated little substitution saturation for this dataset (Xia 2009).

**Fig. 1.**
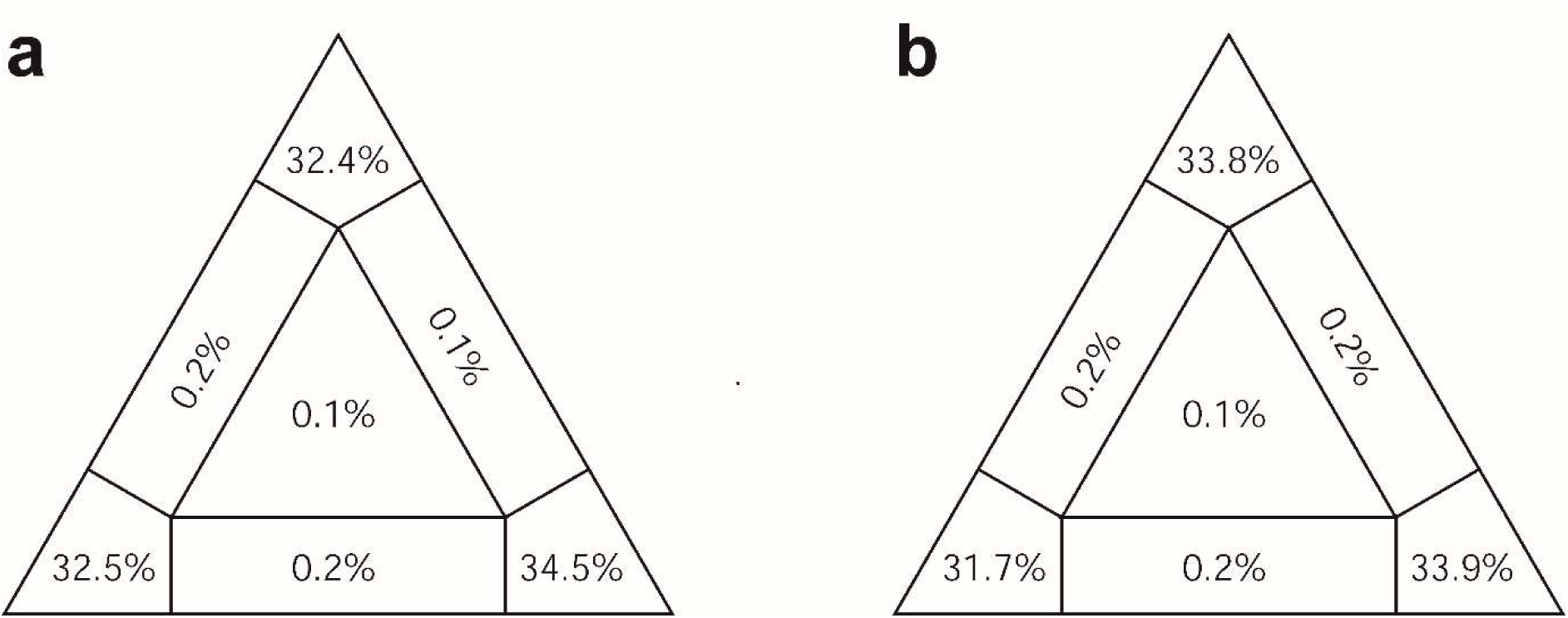
Likelihood mapping (a) before and (b) after Gblocks trimming, demonstrating the high phylogenetic content and suggesting there is no need for alignment cleaning in the case of this dataset.

Using the aligned sequences, two pairwise comparisons between members of the same monogenean family (*C. halli versus C. mbirizei*; *G. nyanzae versus M. karibae*) were made. Firstly, we visualised the nucleotide diversity by a sliding window analysis of nucleotide diversity (π) in DnaSP v. 5.10.01 (Librado & Rozas 2009), with a window size of 300 bp and a step size of 10 bp. To allow comparison between the dactylogyrid and gyrodactylid haplotypes, this approach was limited to the PCGs and rRNA genes. Secondly, for the PCGs of the same pairs of species, the proportion of non-synonymous *versus* synonymous substitutions (dN/dS ratio) was calculated in the codeml program of PAML (Yang 2007) as implemented in PAL2NAL (Suyama et al. 2006).

To situate the African monogeneans under study within their respective families, the PCGs and rRNA genes of all available dactylogyrid (Zhang et al. 2016, 2017) and gyrodactylid (Huyse et al. 2007, 2008; Plaisance et al. 2007; Ye et al. 2014, 2017; Bachmann et al. 2016; Zhang et al. 2016; Zou et al. 2016) mitogenomes were included in phylogenetic analyses. The species of Capsalidae, also monopisthocotylean monogeneans, for which mitogenomes are available (Perkins et al. 2010; Kang et al. 2012; Zhang et al. 2014) were used as outgroup.

The best partition scheme and the optimal models of molecular evolution were determined based on the Bayesian Information Criterion using ModelFinder (Kalyaanamoorthy et al. 2017) with partition merging (Chernomor et al. 2016). The selected partitions and models are shown in Table 1. These were used for Bayesian inference (BI) of phylogeny, whereby posterior probabilities were calculated in MrBayes v. 3.2 (Ronquist et al. 2012) over 10 million generations, sampling the Markov chain at a frequency of 100 generations. Chain stationarity was evidenced by a standard deviation of split frequencies of 8.10^-4^, absence of a trend in the probabilities plotted against the generations, and a potential scale reduction factor (Gelman & Rubin 1992) converging towards 1. One-fourth of the samples were discarded as burn-in. The same partitions were used in a maximum likelihood (ML) search in IQ-TREE (Nguyen et al. 2015), using four gamma-rate categories and an edge-linked partition model. Nodal support was assessed through 10,000 ultrafast bootstrap (Minh et al. 2013) and 1000 Shimodaira-Hasegawa-like approximate likelihood ratio test (Guindon et al. 2010) replicates. In addition, a ML tree was constructed in RAxML v. 8.1.21 (Stamatakis 2006) implemented in raxmlGUI v.1.3 (Silvestro & Michalak 2012), using codon-specific partitions under the GTR + Γ + I model with joint branch length optimization, and with 1000 bootstrap samples to calculate support values. ALTER (Glez-Peña et al. 2010) and GenBank 2 Sequin (Massouh et al. 2016) were used for file conversion, and SequenceMatrix (Vaidya et al. 2011) to concatenate alignment files. Gene orders were compared, and family diagrams of gene orders constructed, using CREx (Bernt et al. 2007). For those genes that were available from the partial mitogenomes, gene order was identical between *M. karibae* and *G. nyanzae*, and between *C. mbirizei* and *C. halli*, respectively. Therefore, only the complete mitogenomes could be included in gene order analyses. For the same reason of comparability, non-coding regions (NCRs) were omitted in gene order analysis.

**Table 1.** Best partition scheme for the dataset of two ribosomal RNA genes and 12 protein-coding genes in the mitochondrial genomes of 14 monopisthocotylean monogenean flatworms. The number of gamma rate categories was set to four.

| Partition | Model of molecular evolution |
| --- | --- |
| 12S + 16S + first codon positions of<br>*atp6, nad2, nad3, nad4, nad4L, nad5, nad6* | TIM2+I+G |
| second codon positions of *atp6, nad1, cox1, cox2, cox3, cytb* | TVM+I+G |
| third codon positions of *atp6, nad1, nad2, nad3, nad4, nad4L, nad5, nad6, cox1, cox2, cox3, cytb* | TIM2+I+G |
| first codon positions of *nad1, cox1, cox2, cox3, cytb* | TIM2+I+G |
| second codon positions of *nad2, nad3, nad4, nad4L, nad5, nad6* | TVM+I+G |

## Results

Genomic DNA sequencing on ¾ of a MiSeq v. 3 flowcell yielded 15,980,972 indexed paired-end 300 bp reads. Complete mitochondrial genomes were assembled for *G. nyanzae* (with a length of 14,885 base pairs (bp)) and *C. halli* (15,047 bp). A circular genome could not be assembled for *C. mbirizei* (12,921 bp) and *M. karibae* (13,002 bp) (Fig. 2). The annotated sequences were deposited in NCBI GenBank under accession numbers xxxxxxxx-x. The total number of reads mapped on the trimmed mt genomes was 12,776, accounting for 0.8 % of the genomic readpool obtained, with an average coverage of 160, 31, 76 and 42 reads and a minimum coverage of 84, 11, 1 and 1 reads for *G. nyanzae*, *C. halli*, *C. mbirizei* and *M. karibae*, respectively. The ribosomal operons of *G. nyanzae* (6,799 bp), *M. karibae* (6,675 bp), *C. halli* (7,496 bp) and *C. mbirizei* (7,005 bp) were deposited as additional molecular vouchers for these species, under NCBI GenBank accession numbers xxxxxxxx-x. For want of complete ribosomal operons for other species targeted in our analyses, these sequences were not considered in our phylogeny.

**Fig. 2.**
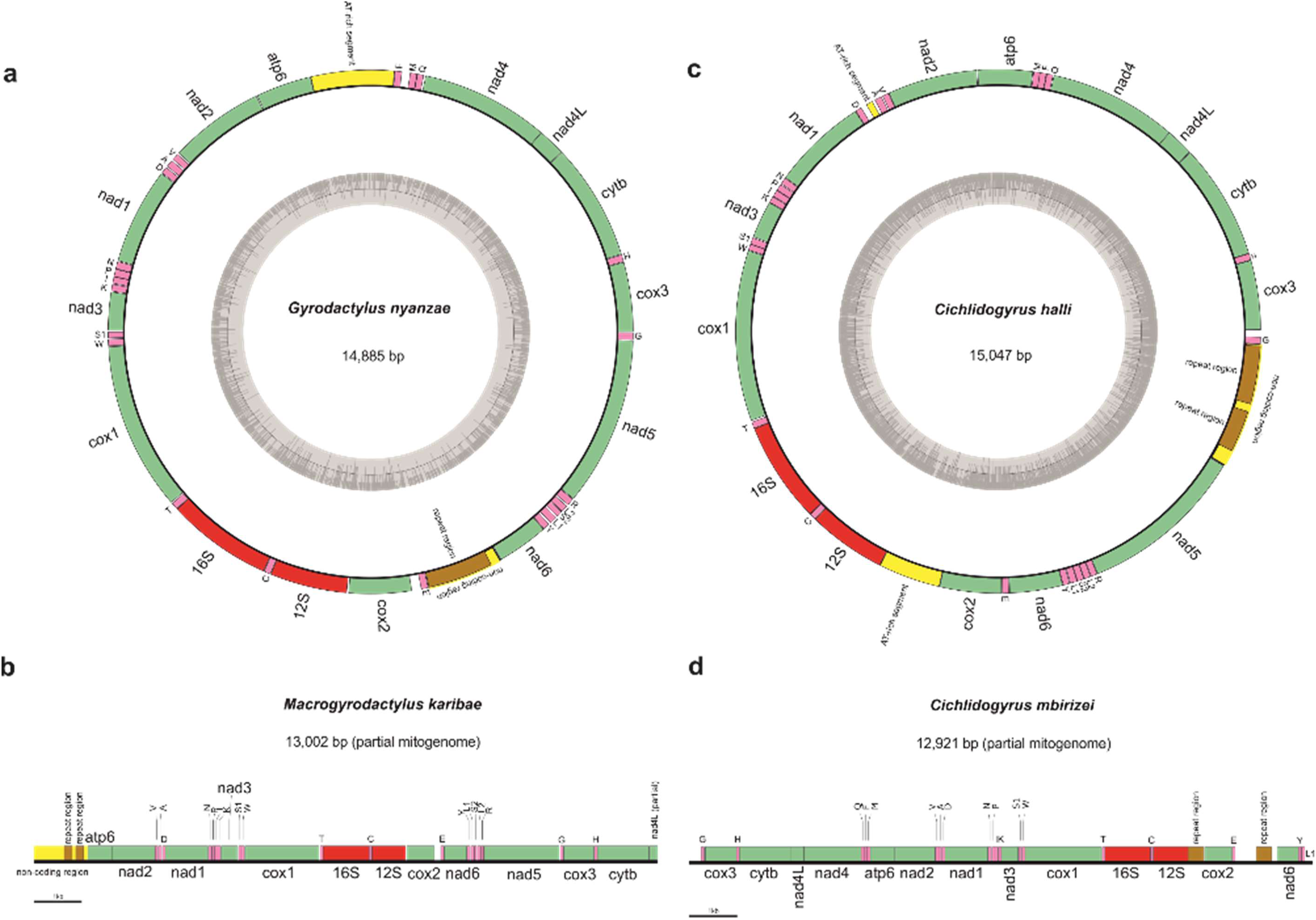
Mitochondrial genomes of four African monogeneans, including two members of the Gyrodactylidae: (a) *Gyrodactylus nyanzae*, (b) *Macrogyrodactylus karibae* (partial genome) and two representatives of the Dactylogyridae: (c) *Cichlidogyrus halli* and (d) *Cichlidogyrus mbirizei* (partial genome). The GC content is displayed for complete mitogenomes.

### Mitogenome characterisation

The protein-coding, ribosomal RNA and tRNA genes are characterised in Table 2. The two complete mitogenomes comprised of 22 tRNA genes (including two for the amino acids serine and leucine each) and 12 intron-free PCGs and lack the *atp*8 gene. The genes coding for the large and small subunit of the mitochondrial rRNA were identified for all four species, as were most PCGs (Fig. 2). Only the *nad*5 gene of *C. mbirizei* and the *nad*4 gene and part of the *nad*4L gene of *M. karibae* were missing. Within the respective monogenean families, start and stop codons of most markers are conserved in these African species. Within the two gyrodactylids, only the stop codons of the *cyt*b, *atp*6, *cox*1 and *nad*6 genes differ; within dactylogyrids, this is only the case for the genes coding for *cyt*b, *nad*3 and *cox*1. The only difference in start codon usage was found in the *nad*2 and *nad*6 gene in *Cichlidogyrus*. Abbreviated stop codons occur in the *cox*3 and *nad*2 genes of the two species of *Cichlidogyrus*.

**Table 2.**
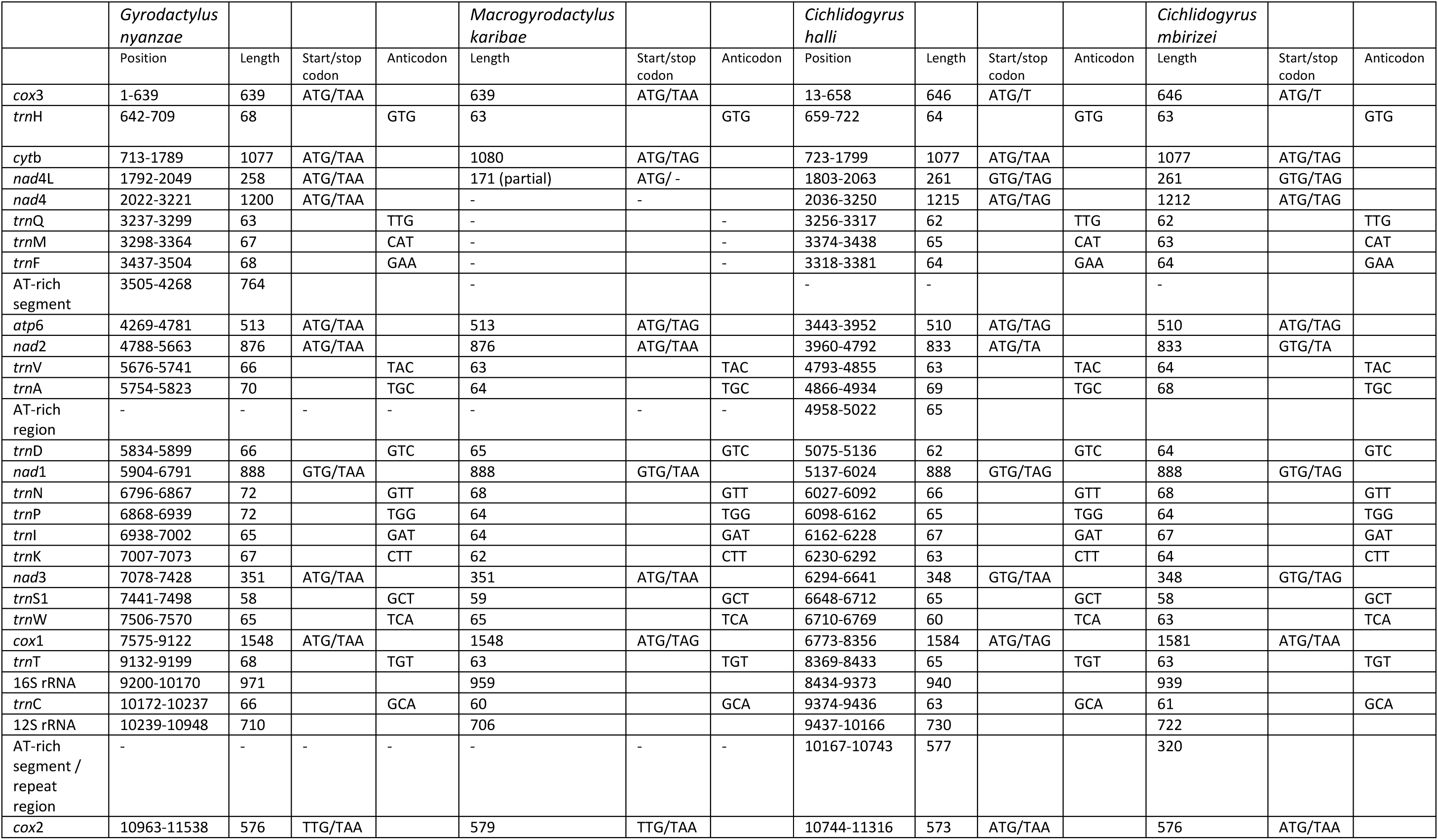
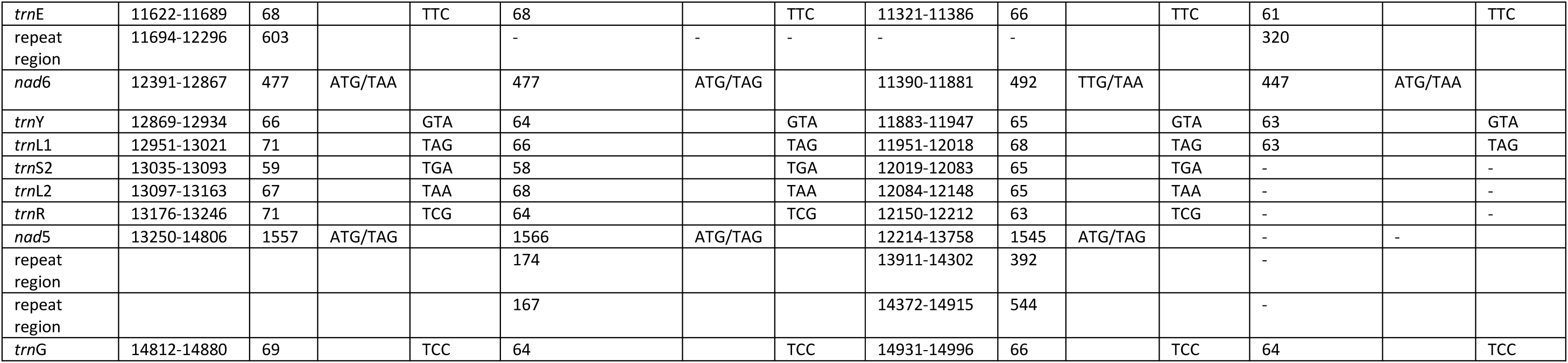
Overview of the length of markers, the start and stop codons (protein-coding genes) and anticodons (tRNA genes) for the four assembled mitochondrial genomes. Positions on the genome are only indicated for complete mitogenomes.

Mitogenome make-up between the African representatives of the dactylogyrids and gyrodactylids only differed in a single tRNA gene transposition. Protein-coding genes appeared in an identical order (see below for pairwise gene order comparisons in a phylogenetic context). Several non-coding regions (NCRs) were observed in all four mitogenomes (Fig. 2). In *G. nyanzae*, one of them, a 603 bp stretch between the genes for *nad*6 and *trn*E, nearly perfectly repeats (except for one substitution) a fragment of 282 bp 2.1 times. The second one, an AT-rich segment (ca. 17% GC content) of 764 bp between the *atp*6 and *trn*F genes, was not identified as a repeat region. In contrast to this, and to the single repeat region of *G. nyanzae*, two consecutive repeat regions were identified adjacent to the *atp*6 gene in the partial mitogenome of *M. karibae*, one 174 bp long with a period of 87 bp (two repeats, 95% match) and the other one 167 bp long with a period of 73 bp (2.3 repeats, 99% match). It has to be noted however, that the possibility of a second non-coding region cannot be excluded due to the double amount of reads in this non-coding region. However, the annotation is incomplete and the exact location can only be inferred using conventional Sanger sequencing. Also the mitogenome of *C. halli* has two repeat regions, between the *trn*G and *nad*5 genes: a 392 bp fragment with repeats of 86 bp (4.6 repeats, 99% match), and a 544 bp fragment with repeats of 167 bp (3.3 repeats, 98% match). In addition, there are AT-rich segments between the *cox*2 and 12S rRNA genes (577 bp with a GC content of ca. 20%) and between the *trn*D and *trn*A genes (65 bp with a GC content of ca. 33%, displaying 58% sequence similarity with a motive in the former AT-rich segment). In the mitogenome of its congener *C. mbirizei*, a 320 bp stretch is duplicated (97% match) between the genes coding for *cox*2 and 12S rRNA on the one hand, and *nad*6 and *trn*E on the other hand.

The sliding window analysis showed concurring patterns and similar values of nucleotide diversity across the mitochondrial genes for the gyrodactylid and dactylogyrid comparisons (Fig. 3). The highest values were found in the genes coding for subunits of NADH dehydrogenase. The dN/dS ratios in the two pairwise comparisons vary, with generally higher values for the genes coding for subunits of NADH dehydrogenase and ATP synthase (Fig. 4). Values remain around or below 0.1 and are higher for the comparison between the two dactylogyrids than between the two gyrodactylids.

**Fig. 3.**
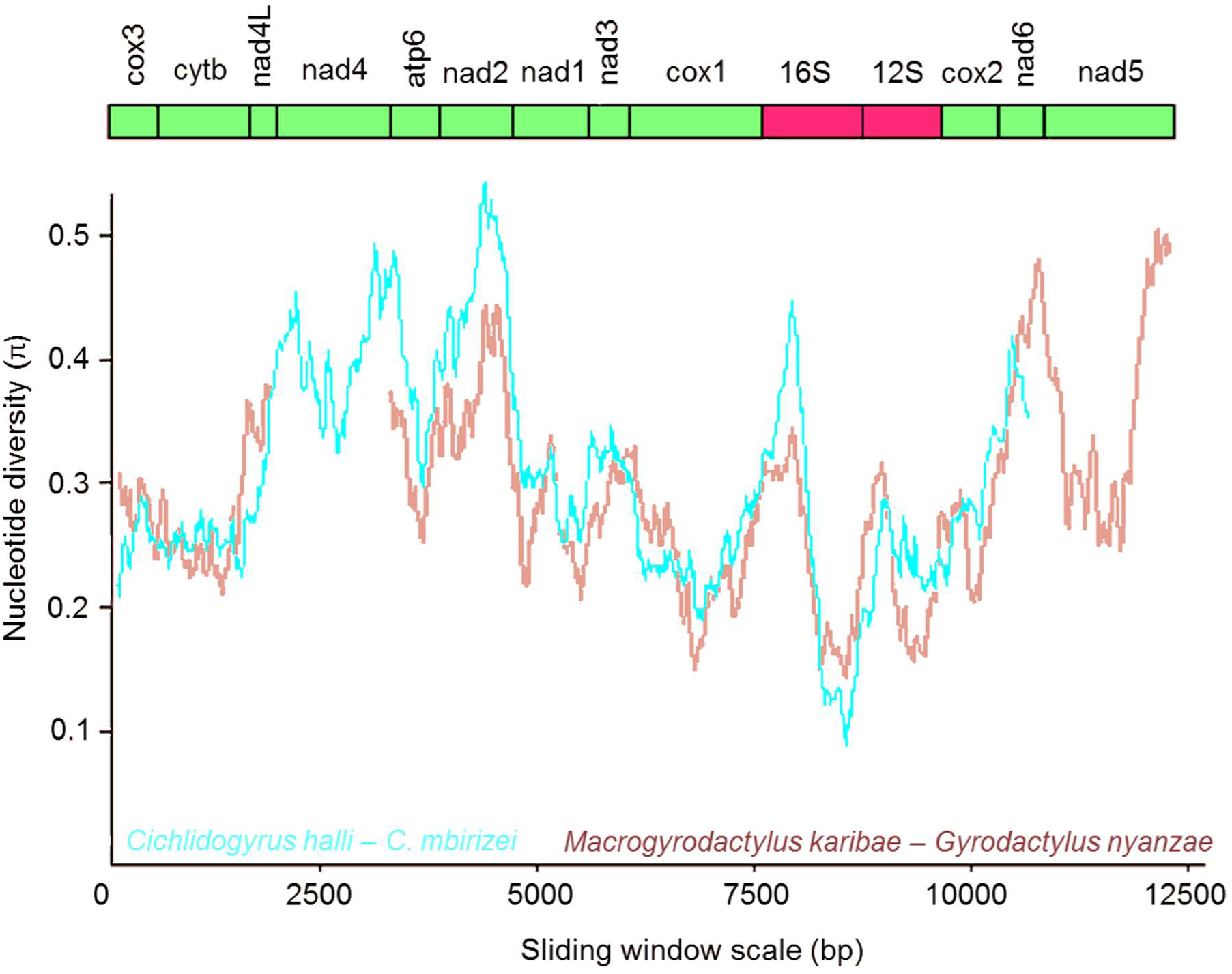
Sliding window analyses (window size 300 bp, step size 10 bp) of the alignment of mitochondrial protein-coding and ribosomal RNA genes used for the phylogenetic analyses of the four mitochondrial genomes of African monogeneans. The lines indicate the nucleotide diversity between two dactylogyrids (*Cichlidogyrus halli* and *C. mbirizei*, in blue) and two gyrodactylids (*Gyrodactylus nyanzae* and *Macrogyrodactylus karibae*, in red). Gene boundaries are indicated above the graph.

**Fig. 4.**
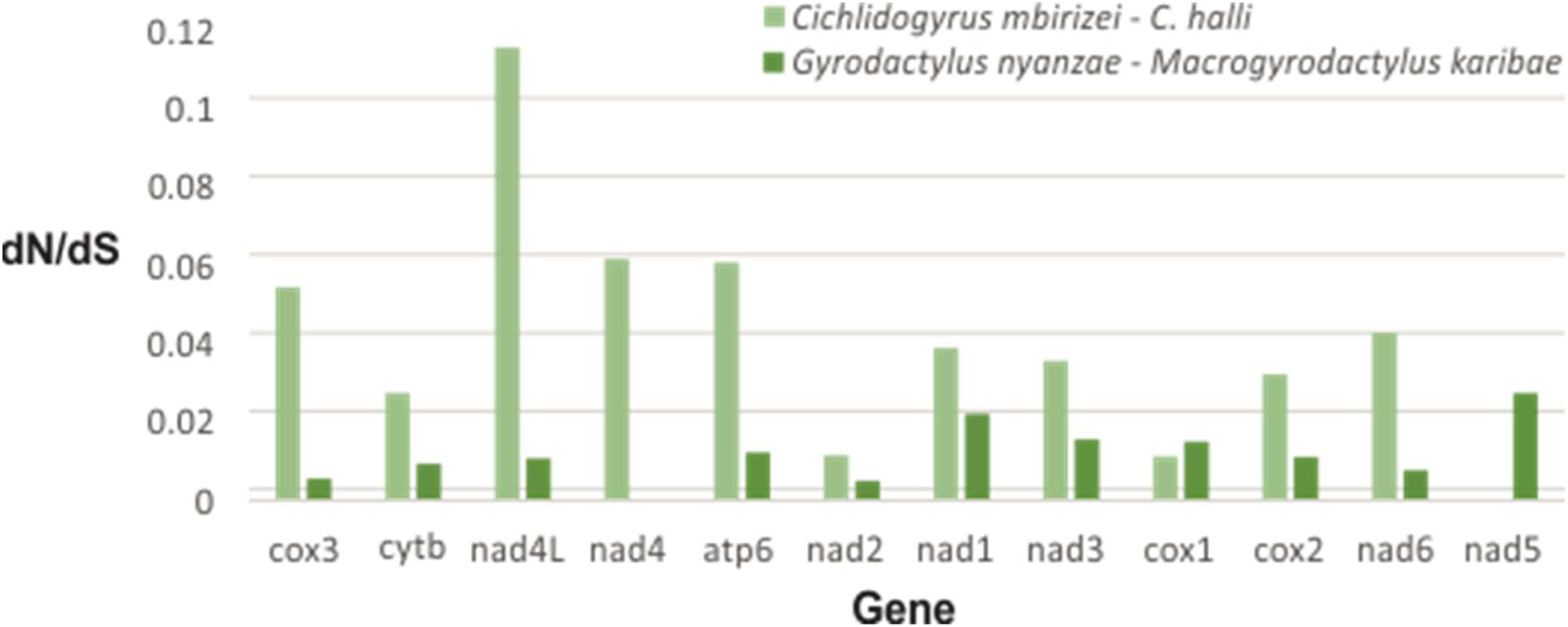
Ratio of non-synonymous to synonymous substitution rates for the protein-coding genes in two pairwise comparisons, between the mitogenomes of African dactylogyrid and gyrodactylid monogeneans, respectively. For *Macrogyrodactylus karibae*, no *nad*4 sequence was available, while the *nad*5 gene was lacking for *Cichlidogyrus mbirizei*.

### Phylogenetic and gene order analyses

The concatenated alignment of 12 PCGs and two rRNA genes for 18 monogenean species contained 12,464 bp and 9,184 variable sites, of which 8,060 were parsimony-informative (although we do not analyse the data with parsimony). The topologies retrieved in ML and BI analyses were near-identical, except for the position of *Tetrancistrum nebulosi*; the resolution within the Dactylogyridae is poor (Fig. 5). Capsalids and dactylogyrids firmly cluster together. *Macrogyrodactylus karibae* and *Paragyrodactylus variegatus* appear as sister taxa, albeit with long branches, presumably due to incomplete taxon coverage. *Gyrodactylus nyanzae* clusters with the clade of *Macrogyrodactylus* and *Paragyrodactylus*, rendering *Gyrodactylus* paraphyletic. *Aglaiogyrodactylus* is firmly positioned as basal to the other gyrodactylids. Within the Gyrodactylidae, a transposition of two tRNA genes was the only difference in gene order between the African representatives and the Palearctic species of *Gyrodactylus* (Fig. 6a), while two adjacent tRNA genes were transposed between the African representatives and *P. variegatus* (Fig. 6b). The difference in mitochondrial gene order between the African gyrodactylids and the Neotropical *Aglaiogyrodacylus forficulatus* can be explained by a tandem duplication random loss (TDRL) event and two transpositions, or, alternatively, four transpositions (Fig. 6c). The gene order in the mitogenomes of both species of *Cichlidogyrus* was identical to that of their family member *T. nebulosi*, and the gyrodactylid *P. variegatus*. This gene order differed simply in one tRNA gene transposition from that of *Gyrodactylus nyanzae* (Fig. 6b) and from that of *Dactylogyrus lamellatus* (Fig. 6d).

**Fig. 5.**
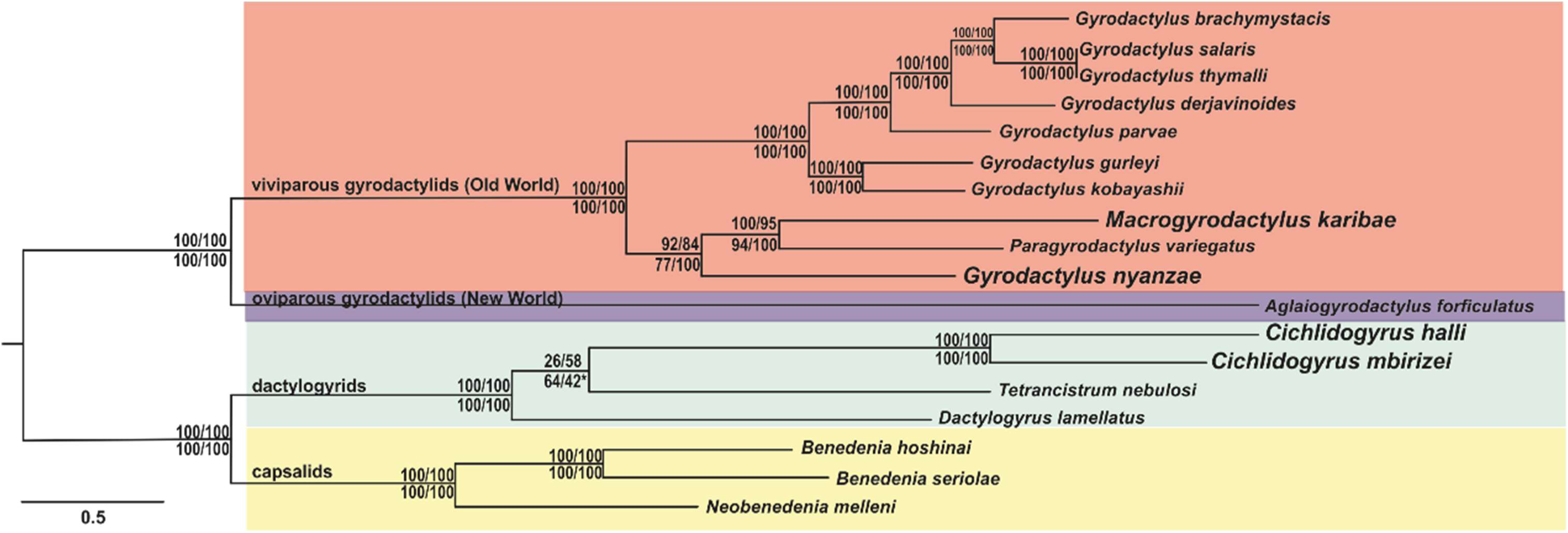
Maximum likelihood phylogram of monopisthocotylean monogeneans based on 12 protein-coding and two ribosomal RNA genes. Support values displayed from (above branch): Shimodaira-Hasegawa-like approximate likelihood ratio test / ultrafast bootstrap, both implemented in IQ-TREE, (below branch) bootstrap in RAxML / Bayesian inference (posterior probability) in MrBayes. An asterisk (*) indicates that this partition was not withheld in the Bayesian consensus tree; the clade grouping *Dactylogyrus lamellatus* and *Tetrancistrum nebulosi* as sister to a monophyletic *Cichlidogyrus* was supported by a posterior probability of 58%. Branch lengths indicate the expected number of substitutions per site.

**Fig. 6.**
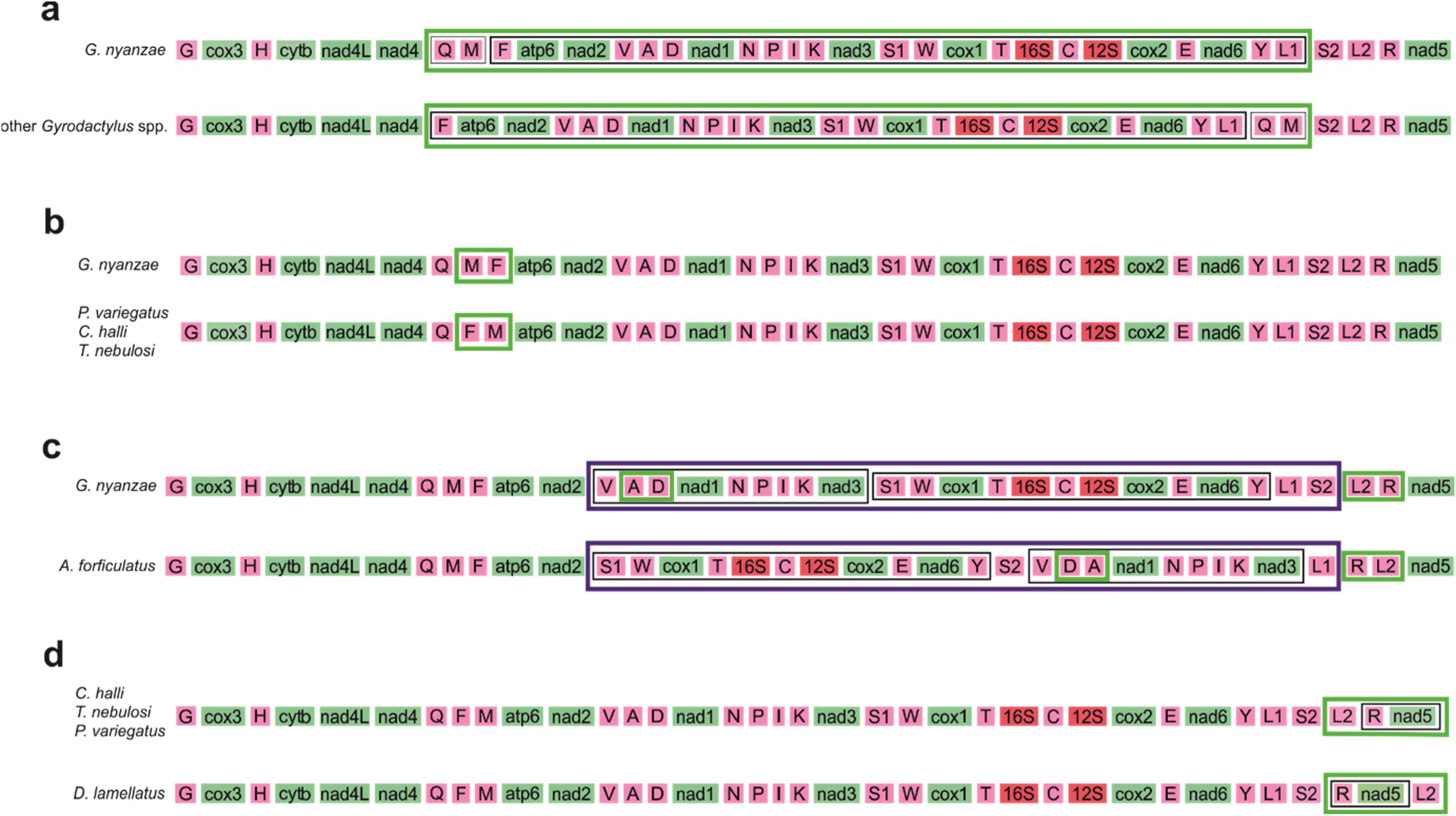
Family diagram explaining gene order changes between (a) African *Gyrodactylus nyanzae* and its Palearctic congeners (a single transposition), (b) *G. nyanzae* and *Paragyrodactylus variegatus* (a single transposition), (c) *G. nyanzae* and the Neotropical oviparous gyrodactylid *Aglaiogyrodactylus forficulatus* (two transpositions and a tandem duplication random loss event (TDRL)) and (d) *Dactylogyrus lamellatus* and the other dactylogyrids (a single transposition). Green boxes indicate transpositions, a dark blue box a TDRL. Only protein-coding genes, tRNA genes and rRNA genes of species for which a complete mitogenome was assembled, are shown.

## Discussion

As the low number of available genetic markers imposes limitations on research on non-model flatworms (Vanhove et al. 2013), improved and cost-efficient NGS offers ever-more opportunities for genomic work on helminths (Wit & Gilleard 2017). Using Illumina technology we assembled, for African gyrodactylid and dactylogyrid monogeneans, one complete and one partial mitogenome each (Fig. 2).

So far only nine gyrodactylid (Huyse et al. 2007, 2008; Plaisance et al. 2007; Ye et al. 2014, 2017; Bachmann et al. 2016; Zhang et al. 2016; Zou et al. 2016) and two dactylogyrid (Zhang et al. 2016, 2017) monogenean mitogenomes have been published. Our study substantially increases the quantity of available mitogenomic data on these two most diverse monogenean families, by one-third, and offers the first mitogenomes from African representatives. The mitochondrial nucleotide diversity of monogeneans is aptly illustrated by the fact that universal barcoding primers for these species-rich helminths are unavailable (Vanhove et al. 2013). Hence utilising High-Throughput Sequencing (HTS) technologies is promising for monogeneans and for other non-model organisms for which typically few or no PCR primers are available. Newly obtained mitogenomes can provide a relatively large set of (coding) molecular markers for molecular evolutionary research. These can be used to develop taxon-specific mitochondrial primers for phylogeographic or population genetic analyses (see below). Challenges however remain, such as the characterisation of AT-rich and repeat regions, in view of the read length of only 300 bp (see also Briscoe et al. 2016a). Also, it is questionable to what extent this NGS approach is workable for rare or opportunistically collected monogenean species, as it has been applied mostly on pools of a considerable number of individuals (this study) or on single larger worm specimens (e.g. Briscoe et al. 2016a). Furthermore, in view of frequent mixed infections, ideally specimens are morphologically identified prior to DNA extraction. This renders the pooling of specimens labour-intensive and sensitive to contamination. Developing reliable NGS shotgun methodologies that can work with single monogeneans, often very small (< 500-1000 µm) in length, will be a worthwhile goal for future molecular ecological and evolutionary studies.

### Mitogenome characterisation and potential for marker development

Throughout the PCGs in the four African mitogenomes, the typical start codons are mostly used: commonly ATG in gyrodactylids, and a combination of ATG and GTG in dactylogyrids. The same goes for the stop codons, typically TAA or TAG. Noteworthy exceptions are the *cox*2 gene of *G. nyanzae* and *M. karibae* and the *nad*6 gene of *C. halli*, with TTG as start codon. This has been reported in monogeneans before, e.g. in the *cox*2 gene of *Paragyrodactylus variegatus* (Ye et al. 2014). However, it is reported for the first time here from a dactylogyrid monogenean (Zhang et al. 2014, 2017); also, it is hitherto unique for a member of *Gyrodactylus*. It is somewhat unsurprising that the full breadth of codon usage diversity in this genus had not yet been captured, since existing mitogenomic data were limited to Palearctic species, all belonging to the subgenus *Limnonephrotus*, defined by Malmberg (1970) on the basis of the excretory system. As regards abbreviated stop codons, the use of T had already been observed in a dactylogyrid monogenean, namely *Dactylogyrus lamellatus* (Zhang et al. 2017). The occurrence of TA as an incomplete stop codon, such as in the *nad*2 gene of both species of *Cichlidogyrus*, is newly reported for dactylogyrids. It has previously been reported in the same gene for *Gyrodactylus brachymystacis* (Ye et al. 2017).

Mitochondrial markers have a wide range of applications in micro-evolutionary and macro-evolutionary research on helminths. For example, for most gyrodactylid and dactylogyrid monogeneans, a small set of established mitochondrial gene fragments (coding for *cox*1, *cox*2, *nad*2 and 16S rRNA) are the most variable markers available. These were applied in population genetics and demography (Bueno-Silva et al. 2011; Kmentová et al. 2016), in barcoding (Hansen et al. 2007; Bueno-Silva et al. 2014), in phylogeography (Meinilä et al. 2004; Plaisance et al. 2008; Wang et al. 2016; Huyse et al. 2017) and to detect hybridisation (Barson et al. 2010). Mitochondrial sequences also served to elucidate the phylogeny of closely related species (Vanhove et al. 2015), genera (Plaisance et al. 2005; Bachmann et al. 2016) or higher-order taxa in monogeneans (Park et al. 2007) and other flatworms (e.g. tapeworms: Waeschenbach et al. 2012). The availability of mitogenomic information on African monogeneans may facilitate the development of a wider range of mitochondrial markers applicable on this understudied tropical fish parasite fauna.

Within Palearctic gyrodactylids, *nad*2, *nad*4 and *nad*5 are the most variable genes in the mitochondrial genome and were therefore suggested as markers to study population-level processes such as transmission and host-switching, and to detect pathogenic strains (Huyse et al. 2008; Ye et al. 2017). Insofar as these genes were available in our alignment, this pattern is confirmed for African gyrodactylids and dactylogyrids; especially the *nad*2 gene seems promising for marker development as it is flanked by rather conservative stretches (Fig. 3). In light of massive anthropogenic translocations of African cichlids and clariids, we may add the detection of introduced monogenean strains as a potential application of these markers. These more variable markers roughly correspond with those displaying a higher dN/dS ratio (Fig. 4); hence, those genes are under a somewhat relaxed selection. This, and the fact that dN/dS values for all mitochondrial PCGs fall well below 1, which indicates purifying selection, confirms earlier mitogenomic work on monogeneans (e.g. Huyse et al. 2008; Zhang et al. 2017). Overall purifying selection acting on mitochondrial genes, albeit more relaxed for e.g. certain NADH dehydrogenase complex genes, has also been observed in a range of vertebrates (Castellana et al. 2011; Jakobsen et al. 2016).

All hitherto known mitogenomes of species of *Gyrodactylus*, all representing the subgenus *Limnonephrotus*, contain two near-identical NCRs (Ye et al. 2017). Conversely, such duplicated NCRs are absent in their congener *G. nyanzae* and, in our dataset, only found in *C. mbirizei*. Indeed, our results suggest substantial differences in the length, number and position of NCRs between African monogeneans even among gyrodactylids and within *Cichlidogyrus* (Fig. 2). There is no clear phylogenetic pattern, but a comparison with mitochondrial genomes of other gyrodactylid and dactylogyrid monogeneans indicates that non-coding (repeat) regions are commonly positioned in between certain pairs of genes: e.g. *trn*D and *trn*A in *C. halli* and *Aglaiogyrodactylus forficulatus* (Bachmann et al. 2016); *trn*E and *nad*6 in *G. nyanzae* and *C. mbirizei*; *nad*5 and *trn*G in *C. halli* and *Tetrancistrum nebulosi* (Zhang et al. 2014); *trn*F and *atp*6 in *G. nyanzae* and its Palearctic congeners (e.g. Huyse et al. 2008; Ye et al. 2017); and 12S rRNA and *cox*2 in *C. halli* and *C. mbirizei*. Also a NCR containing two repeat regions in the vicinity of the *nad*5 gene, such as reported here for *C. halli*, has been reported before in dactylogyrids, namely by Zhang et al. (2017) for *Dactylogyrus lamellatus*. We reiterate these authors’ suggestion about the potential these tandem repeats offer for population-level research on monogeneans. The possibility of a functional role of these AT-rich and repeat regions has already been raised in multiple previous studies (e.g. Ye et al. 2014). The similarities we found in the position of these NCRs throughout a range of dactylogyrid and gyrodactylid mitogenomes may add further proof to the significance of the location of these regions throughout monopisthocotylean monogeneans.

### Ribosomal operons and utility

Characterising full nuclear ribosomal operons provides a wealth of information for established and prospective molecular markers. Ribosomal DNA codes for all the nuclear ribosomal genes (ETS, 18S, ITS1, 5.8S, ITS2 and 28S) including the external and internal transcribed spacer regions (ETS, ITS respectively). As tandemly repeated units, ribosomal operons occur in high number, and the remarkable variation in rate of molecular evolution within and between nuclear rRNA gene regions has driven their popularity as a source for molecular markers in Metazoa (Mallatt et al. 2010) and within the parasitic flatworms (Lockyer et al. 2003). Within flatworms ITS regions are popular for discriminating between closely related species (Nolan & Cribb 2005), and numerous authors have used complete 18S and partial (D1-D3) regions of 28S rDNA for phylogenetics of Monogenea (e.g. Olson & Littlewood 2002). In combination with mitochondrial genes, nuclear ribosomal RNA genes are invaluable for discriminating hybrid species, especially important when revealing the identity of disease-causing parasites (e.g. Huyse et al. 2013). Many nuclear rRNA gene regions have been used to discriminate species, to resolve phylogenetic relations and as molecular ecological markers amongst monogeneans (Mendlová et al. 2012; Vanhove et al. 2015). Within the newly characterised mitogenomes of African monogeneans in this study, the full operon ranged in size between 6,675-7,496 bp largely reflecting differences in length of spacer regions. We consider this to be a rich resource for a diversity of future studies, especially in the emerging field of environmental (eDNA) metabarcoding and metagenomics where access to highly conserved, and high copy number, markers will greatly benefit accurate species identification (Tang et al. 2012). In addition, a pairwise or multiple alignment of full ribosomal operons will readily highlight regions of sequence variability and conservation suggesting potential markers for a variety of purposes and regions of conservation potentially useful for PCR primer design. Future studies aimed at population genetics, hybridisation, biogeography, cryptic species recognition, and host-parasite interactions will benefit from access to the full rRNA operon and the full mitogenomes of these, and additional taxa. Certainly, characterisation of full ribosomal operons by means of NGS genome skimming is considerably easier, and cheaper than by long PCR and primer walking using Sanger technology.

### Mitochondrial phylogeny, gene order and implications for the position of African gyrodactylid and dactylogyrid monogeneans

Our phylogenetic reconstruction based on 12 mitochondrial PCGs and 2 rRNA genes aimed to elucidate the position of African *Macrogyrodactylus*, *Gyrodactylus* and *Cichlidogyrus* (Fig. 5). All tree topologies firmly place the Neotropical oviparous *Aglaiaogyrodactylus forficulatus* as a sister lineage to all other representatives of the Gyrodactylidae. This refutes Malmberg’s (1998) hypothesis of *Macrogyrodactylus* being the most basal gyrodactylid. In addition, the inclusion of an African representative, *G. nyanzae*, renders *Gyrodactylus* paraphyletic. Hence, we provide the first mitochondrial data supporting the paraphyly of the genus, corroborating earlier phylogenetic hypotheses based on morphology (Kritsky & Boeger 2003) or nuclear rRNA genes (Vanhove et al. 2011; Gilmore et al. 2012; Přikrylová et al. 2013).

The relationships between the only three dactylogyrid genera in the mitogenomic tree, all of them from the ‘Old World’, are not well resolved. Both *Cichlidogyrus* and *Tetrancistrum* have previously been mentioned as members of the Ancyrocephalinae (or Ancyrocephalidae). The monophyly of this (sub)family has often been challenged in earlier work (e.g. Šimková et al. 2003, 2006; Plaisance et al. 2005). Two topologies (*Tetrancistrum* as a sister to *Cichlidogyrus* or, alternatively, to *Dactylogyrus*) have an equally low posterior probability under BI. Hence, our tree is not informative on the status of the Ancyrocephalinae *versus* the Dactylogyrinae, to which *Dactylogyrus* belongs. Although the polytomy makes it hard to favour either of the two alternative positions of *Tetrancistrum*, the gene order is identical between the representatives of *Tetrancistrum* and *Cichlidogyrus* in contrast to the representative of *Dactylogyrus*. We therefore consider the sister-group relation between the latter two genera the biologically most likely hypothesis. This also corresponds to the nuclear rDNA-based results of Blasco-Costa et al. (2012) suggesting that *Tetrancistrum* and *Cichlidogyrus* belong to the same clade of mostly marine ancyrocephalines. The affinity between *Cichlidogyrus* and marine genera, despite the likely sampling bias as many dactylogyrid genera have not yet been sequenced, is worth looking into because of the potential of cichlid parasites in elucidating the alleged role of marine dispersal in cichlid biogeography (Pariselle et al. 2011). It would be worthwhile to consider mitochondrial gene order as a phylogenetic marker for further disentangling the relationships between purported dactylogyridean (sub)families.

While it is well-established that gene order is phylogenetically informative, it mainly seems to differ, certainly for PCGs, at the level of major flatworm lineages, such as between catenulids, triclads, polyclads and neodermatans (Rosa et al. 2017). Within the major flatworm clades, e.g. at order or family level, differences in mitogenome architecture mainly concern tRNA genes and NCRs (e.g. Zhang et al. 2014 for capsalids; Solà et al. 2015 for triclads). This is confirmed in our results, where gene order differences within the dactylogyrids (Fig. 6d) and the viviparous gyrodactylids only concern tRNA genes (Fig. 6a,b). The transpositions seem to concur with evolutionary distance, e.g. simply two adjacent tRNA genes have swapped position within the *Macrogyrodactylus-Paragyrodactylus-Gyrodactylus nyanzae* clade (Fig. 6b). All viviparous gyrodactylids including *Macrogyrodactylus* (Fig. 6c) have identical PCG orders. However, *Aglaiogyrodactylus forficulatus* displays a different PCG arrangement, which underscores its particular position within the Gyrodactylidae, apart from the viviparous members of this family.

## Conclusions

The first mitogenomic data for African monogeneans are provided, characterising two partial and two complete mitochondrial genomes. These confirm earlier results on the variability and selection pressures on mitochondrial genes in monogeneans, and highlight some patterns in the location of NCRs. These mitogenomes increased the known diversity of start and stop codon usage in dactylogyrids and in species of *Gyrodactylus*. A phylogeny based on 14 mitochondrial markers firmly confirmed the Neotropical oviparous *Aglaiogyrodactylus* as ‘basal’ to the other gyrodactylids, rather than the allegedly ‘primitive’ *Macrogyrodactylus*. Furthermore, it provided additional evidence for the paraphyly of *Gyrodactylus*. While the gene order for PCGs remained constant throughout the species considered, the study suggested tRNA transpositions to be phylogenetically informative for relationships within the family level.

As highlighted above, (mitochondrial) gene sequences are established tools in the identification of monogeneans, including potentially pathogenic and invasive strains of fish parasites, but their availability for African species remains limited. We hope that this study will contribute to marker development and diagnostics, and hence to ecological and evolutionary studies of African monogeneans.

## List of abbreviations

bp: base pairs
BI: Bayesian inference
HTS: High-Throughput Sequencing
MITObim: mitochondrial baiting and iterative mapping
ML: maximum likelihood
NCR: non-coding region
NGS: next-generation sequencing
PCG: protein-coding gene
rDNA: ribosomal DNA
rRNA: ribosomal RNA
TDRL: tandem duplication random loss event

## Declarations

### Consent to participate

Not applicable

### Ethics approval

In the absence of relevant animal welfare regulations in the D.R. Congo, the same strict codes of practice enforced within the European Union were applied. Sampling was carried out under attestation de recherche no. 863/2014 from the Faculté des Sciences Agronomiques of the Université de Lubumbashi.

### Consent for publication

Not applicable

### Availability of data and materials

The sequence data produced and analysed during the current study were deposited in NCBI GenBank under accession numbers xxxxxxxx-x and xxxxxxxx-x. Voucher specimens are available in the invertebrate collection of the Royal Museum for Central Africa (RMCA), Tervuren, Belgium. The posterior ends (with opisthaptor) of four of the specimens of *Macrogyrodactylus karibae* used were deposited under accession numbers MRAC xxxxx-xx; for the other monogenean species, entire animals were used for DNA extraction, and conspecifics from the same host specimen are available under accession numbers MRAC xxxxx-xx.

### Authors’ contributions

MPMV conceived the study, collected and identified specimens, analysed data and drafted the manuscript. AGB carried out experiments and analysed data. MWPJ prepared and identified specimens. DTJL analysed data, oversaw the study and provided lab facilities. TH conceived and oversaw the study, carried out experiments, analysed data and provided lab facilities. All authors contributed to drafting the manuscript and read and approved the final version of the manuscript.

## Acknowledgements

The authors cordially thank W. Fannes, E.J. Vreven, A. Chocha Manda, E. Abwe, B. Katemo Manda, G. Kapepula Kasembele, M. Kasongo Ilunga Kayaba, C. Kalombo Kabalika, C. Mukweze Mulelenu, M. Katumbi Chapwe and J. Snoeks for their help with sampling and/or planning this research, I. Přikrylová and A. Pariselle for advice on monogenean identification, and R. Väinölä and M. Monnens for helpful input regarding the analyses.

